# Contribution of unfixed transposable element insertions to human regulatory variation

**DOI:** 10.1101/792937

**Authors:** Clément Goubert, Nicolas Arce Zevallos, Cédric Feschotte

## Abstract

Thousands of unfixed transposable element (TE) insertions segregate in the human population, but little is known about their impact on genome function. Recently, a few studies associated polymorphic TE insertions to mRNA levels of adjacent genes, but the biological significance of these associations, their replicability across cell types, and the mechanisms by which they may regulate genes remain largely unknown. Here we performed a TE-expression QTL analysis of 444 lymphoblastoid cell lines and 294 induced pluripotent stem cells using a newly developed set of genotypes for 2,806 polymorphic TE insertions. We identified 211 and 324 TE-eQTL acting in *cis* in each respective cell type. Approximately one fourth were shared across cell types with strongly correlated effects. Furthermore, analysis of chromatin accessibility QTL in a subset of the lymphoblastoid cell lines suggests that unfixed TEs often modulate the activity of enhancers and other distal regulatory DNA elements, which tend to lose accessibility when a TE inserts within them. We also document a case of an unfixed TE likely influencing gene expression at the post-transcriptional level. Our study points to broad and diverse *cis*-regulatory effects of unfixed TEs in the human population and underscores their plausible contribution to phenotypic variation.

## Background

Transposable elements (TEs) are ubiquitous genetic entities that relocate and multiply within genomes. TE sequences occupy a large fraction of the eukaryotic nuclear DNA, including humans where they account for more than half of the genetic material [1]. New TE insertions represent an important source of structural genomic variation that can affect both coding and regulatory components of the genome[2–4]. Notably, many TEs deposit *cis*-regulatory sequences that can modulate flanking gene expression, and sometimes being repurposed for beneficial cellular function[5–7].

While the human genome hosts hundreds of different TE families from various classes, only three retrotransposon families, LINE1 (L1), Alu and SVA, are known to be active and produce *de novo* insertions, including ∼100 disease-causing insertions thus far documented[8–10]. These three families represent the primary source of unfixed TEs across humans[11, 12]. To date, whole genome sequencing studies have discovered more than 19,000 TE loci segregating in the human population[11–15]. Despite their potential role in shaping human phenotypic variation, including disease susceptibility[16, 17], very little is known about the impact of these polymorphic insertions on genome function. Only a handful of recent studies have started to unravel their contribution to gene expression variation[16–19].

The regulation of gene expression is central to cellular function and differentiation in development and physiology and changes in gene expression are important drivers of phenotypic variation[20]. Steady state mRNA levels partially govern gene expression in response to inputs integrated by various regulatory sequences acting in *cis* or *trans*[20, 21]. Expression quantitative trait loci (eQTL) studies offer a systematic approach to identify such *cis*-regulatory elements by correlating the genotypes of genomic variants segregating in individuals with mRNA levels of specific genes, which can be measured on a large scale by RNA-sequencing (RNA-seq) [22, 23]. Previous eQTL studies have established that virtually every gene in the human genome has its expression affected by at least one genomic variant (typically a single nucleotide polymorphism, SNP) located in *cis* (usually defined as within a maximum distance of 1 Mb) and generally in non-coding regions[21,24–26]. A recent analysis of 44 different tissues by the GTEx consortium indicates that such *cis*-eQTL fall into two broad categories: those shared across most tissues and those apparently acting in a single or a restricted number of similar tissues[21]. Importantly, most eQTL studies thus far have focused on SNPs, yet other type of genomic variants, such as unfixed TEs insertions, are common in the human population and likely to have more drastic effects on gene expression[27–30].

In a pioneering study, Wang et al. [18] mapped TE-eQTL in a reference set of 445 EBV-transformed lymphoblasts (lymphoblastoid cell lines, LCL) for which TE insertion genotypes [12] and RNA-seq data [24] had been previously generated as part of the 1000 Genomes Project and GEUVADIS consortium, respectively. In this dataset, they identified 83 *cis* TE-eQTL where the genotype of TE insertions correlated with mRNA levels of adjacent genes [18]. These data, as well as a follow-up study building on these results[17] suggest that unfixed TEs represent a class of structural variants that plays an important role in driving population- and tissue-specific regulatory variation. However, many questions remain unexplored, in particular (i) the size, direction and strength of the discovered associations, (ii) their tissue- or cell type-specificity and (iii) a better understanding of the molecular mechanisms by which unfixed TEs may modulate gene expression.

To begin filling these gaps, we use a newly assembled set of TE genotypes [31], including more than 800 TE insertions not considered in previous studies [17,18,32], in conjunction with reprocessed RNA-seq quantifications to map TE-eQTL in the aforementioned LCL from 444 individuals. We examined the cell type-specificity of these associations by performing another TE-eQTL mapping in 294 induced human pluripotent stem cells (iPSC) available through the HipSci consortium[24, www.hipsci.org]. To investigate *cis*-regulatory mechanisms by which unfixed TEs may affect adjacent gene expression, we explored their association with chromatin accessibility QTL mapped for a subset of the LCLs. Lastly, we document a case of an unfixed TE insertion likely affecting the expression of a gene involved in lipid metabolism through a post-transcriptional mechanism.

## Results

### Mapping *cis*-TE-eQTL in LCL and iPSC

We conducted a search for TE-eQTL in 444 lymphoblastoid cell lines (LCL) and 294 human induced pluripotent stem cells (iPSC) using the genotypes of unfixed TE insertions. In LCL, the genomic coordinates for these loci (13,986 Alu; 3,104 L1 and 844 SVA) were obtained from the previous analysis of 2,504 human samples from the 1000 Genomes Project [12]. To improve genotyping quality, we re-analyzed the 445 LCL samples with TypeTE [31]. These recalls enabled us to use 860 TE insertions present in the reference genome (hg19), which have not been interrogated in previous studies [17,18,32] possibly due to the uncertainty of their original genotypes [31]. To call TE genotypes in iPSC, we initially used the genomic alignments generated by the HipSci consortium (www.hipsci.org) for 326 healthy fibroblast-derived iPSCs, predominantly sourced from subjects of British ancestry [33]. TE detection and genotyping in this iPSC dataset was achieved using MELT2 [15] and TypeTE (see Methods). We identified 8,477 TE (6,931 Alu; 1,068 L1 and 478 SVA) with a ‘PASS’ flag following MELT2 analysis. These numbers are in line with those expected based on the results of the 1000 Genome Project [12, 15]. After sample filtering (see below), we identified a total of 2,806 unfixed TEs segregating in both 294 iPSC and 444 LCL with a minimum allele frequency of 5% in each dataset.

In order to identify TE**-**expression QTL (TE-eQTL), we first quantified the steady-state RNA levels of LCLs by applying the program Kallisto (version 0.46.0 [34]) against RefSeq (GRCh37.75) using RNA-seq reads originally generated by Lappalainen et al. [35]. This allowed us to match the RNA-seq quantification data generated for the iPSCs by the HipSci consortium (Helena Kilpinen, personal communication; see also Methods). We then analyzed independently LCL and iPSC datasets to search for correlations between TE genotypes and normalized mRNA levels using QTLtools (version 1.1[36]). *cis-*TE-eQTL were searched within a 1 Mb window of associated genes (eGenes). Statistical significance was assessed by performing 10,000 permutations of the gene expression matrix and applying a 5% false discovery rate for multiple testing correction (see Methods)[36]. The results showed that two chromosomal regions produced an exceptionally high density of predicted TE-eQTLs: one corresponds to the HLA locus at 6p21 (chr6: 28,477,797-33,448,354 [hg19]) and the other corresponds to the 17q21.31 inversion (chr17: 38,100,001-50,200,000 [hg19]). We suspect both regions to be prone to yield a high rate of false positives in eQTL analyses due to their complex, highly repetitive nature which makes short read mapping unreliable and hinders both TE insertion mapping/genotyping and RNA-seq quantification[37, 38]. Thus, conservatively, we excluded 49 (LCL) and 53 (iPSC) TE-eQTLs falling within these two regions from subsequent analysis. After this filtering, we obtained a total of 211 and 340 TE-eQTLs in LCL and iPSC, respectively (Figure 1, Table 1 and Figure S.1). Repeating the analysis with increasingly larger, randomly selected subsamples revealed no evidence of saturation regarding the total number of TE-eQTL discovered (Figure S.2).

**Figure 1.**
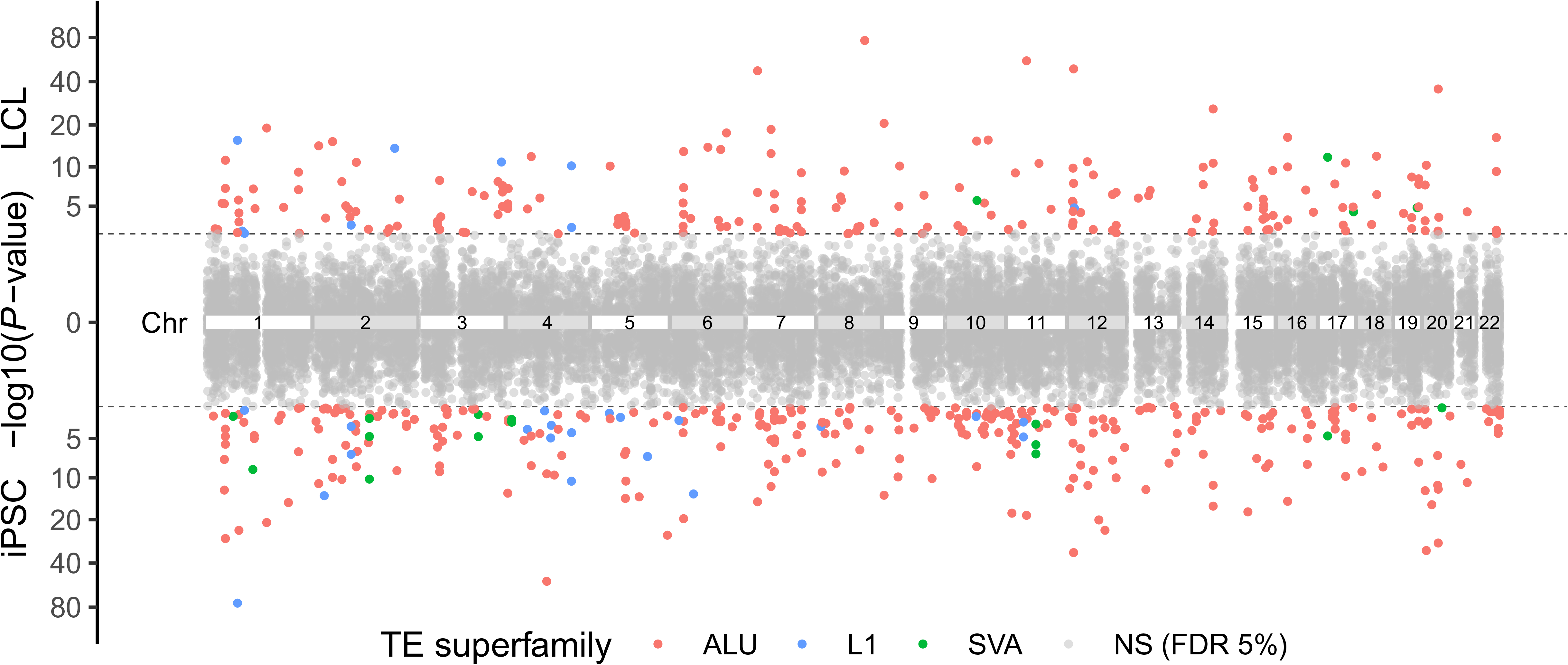
Manhattan plots. P-values distribution according to chromosomal position of TE-eQTL mapped in LCL (up) and in iPSC (down) TE-eQTL. The gray dashed line represents the 5% FDR cutoff (LCL: 1.21×10^-3^ iPSC: 2.18×10^-3^). Each *P*-value is colored according to the TE family.

**Table 1.** TE-QTL in LCL and iPSCs. ‘Genes 50%’: number of genes expressed in at least 50% of the samples and considered in the analysis.

| Cell type | TE-eQTL |  |  |  | TE-caQTL |  |  |  |
| --- | --- | --- | --- | --- | --- | --- | --- | --- |
|  | Sample size | Genes 50% | eGenes | eTEs (Alu/L1/SVA) | Sample size | ATAC Peaks | caPeaks | caTEs (Alu/L1/SVA) |
| LCL | 444 | 14,320 | 211 | 157 / 9 / 4 | 86 | 277,128 | 656 | 394 / 28 / 9 |
| iPSC | 294 | 15,725 | 340 | 242 / 19 / 8 | - | - | - | - |

For both cell type, the vast majority of TEs associated with eQTLs (eTEs) are located within 250 kb of the gene body (From TSS to the end of the 3’-UTR, Figure 2-A,B). To further examine whether the distribution of eTEs relative to eGenes departs from that of all unfixed TEs found within 1 Mb of a quantified gene (Figure 2-C “TE/gene < 1 Mb”), we compared the location of the two TE categories (eTEs and all TEs in a 1Mb window) with randomly sampled genomic positions (see Methods). We found that eTEs are enriched within introns and within a 10-kb window upstream and downstream of eGenes, but depleted in intergenic regions, coding exons and UTRs. By contrast, when all TEs present in the same 1-Mb window are considered, we observe that they are depleted in exons, UTRs, and within 10 kb upstream and downstream regions of genes, but their density in introns and intergenic regions do not depart from the random expectations (Figure 2-C). We conclude that eTEs are more closely associated with eGenes than other TEs, but remain generally excluded from exons and UTRs.

**Figure 2.**
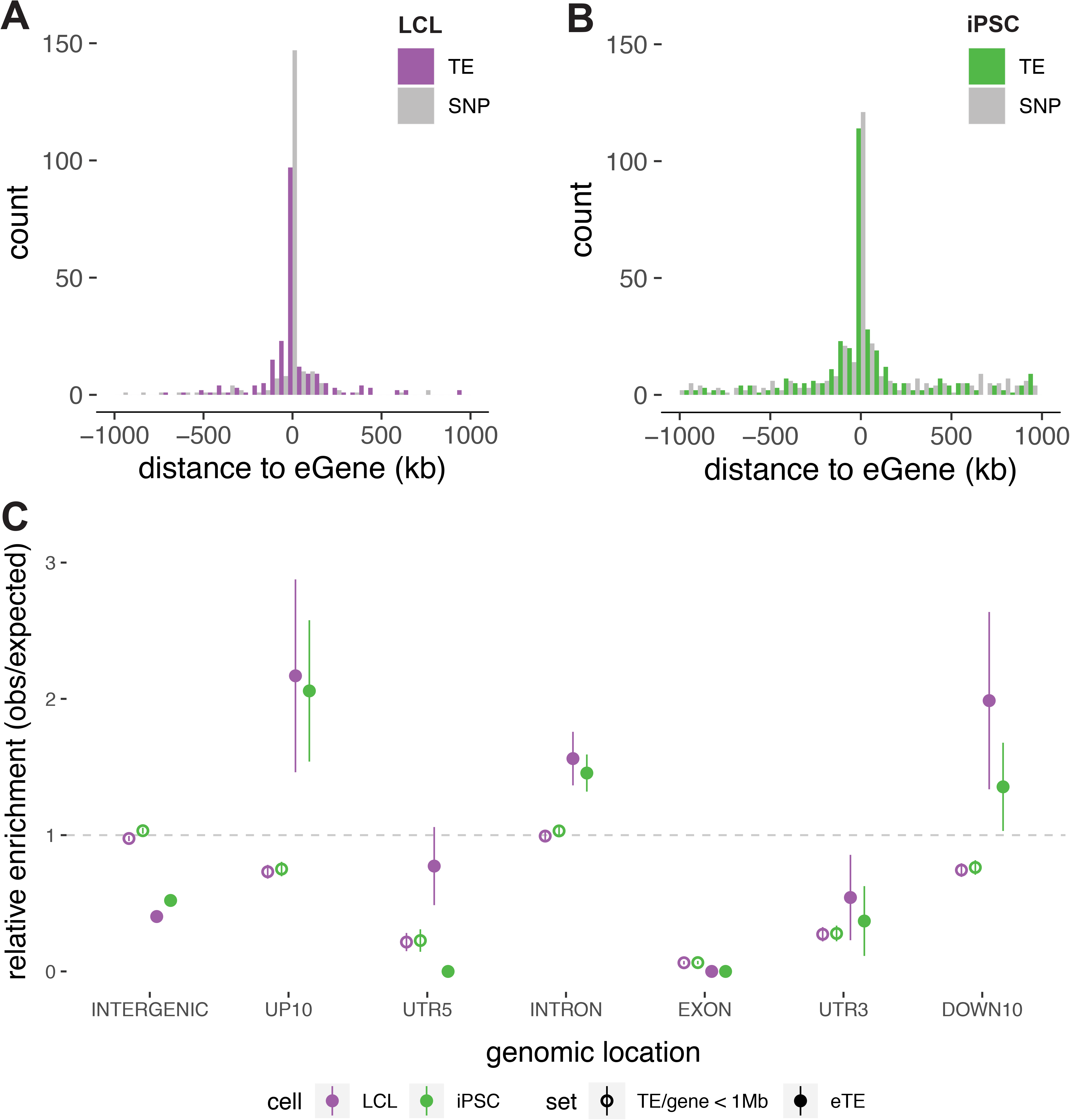
TE-eQTL distribution. A-B - Distribution of the distances between eTE and eGene for LCL (left) and iPSC (right). Grey shading corresponds to the distribution for a matched number of SNP-eQTL. C - TE enrichments in different genomic compartments: expected values are estimated by selecting a random set of genomic intervals equal to the number of each type of variant. TE/gene < 1MB: unfixed TE located within 1Mb of a gene; eTE: TE associated in a given TE-eQTL. UP10: region spanning 10 kb upstream of gene transcription start site; DOWN10: region spanning 10 kb downstream of gene termination site. UTR5: 5’-UTR; UTR3: 3’-UTR. INTERGENIC: genomic region distant > 10 kb upstream TSS or > 10 kb downstream 3’-UTR

### Effect size and TE-eQTLs significance

Next, we focused on the effect size and *P*-value distributions of TE-eQTL as a first step to assess their potential biological relevance. The effect size captures the magnitude of changes in gene expression between the three possible TE genotypes (0/0: no TE, 0/1: heterozygous TE insertion, 1/1 homozygous TE insertion) as the slope of the linear regression. The *P*-value associated to each TE-eQTL reflects the strength of this correlation[39].

We found little to no correlation between the distance of eTEs to eGenes and the effect size of the TE-eQTL (Figure S.3, Spearman correlation test, r = −0.02, *P* > 0.05 for LCL; r = −0.17, *P* < 0.01 for iPSC). Similarly, there was no strong correlation between the distance of eTEs to eGenes and the strength (*P*-value) of the eQTL (Figure S.4, Spearman correlation test, LCL: r = −0.11 *P* > 0.05; iPSC: r = −0.18, *P* < 0.01). However, as expected, effect sizes and *P*-values were positively correlated (Figure S.5, Spearman correlation test, LCL: r = 0.80 *P* < 0.01; iPSC: r = 0.52, *P* < 0.01), which means that there is generally stronger statistical support for TE-eQTL candidates associated with the largest change in gene expression between genotypes.

When we examined TE-eQTL for the different families of TEs (Alu, L1, SVA), we observe distinct trends in the direction of their effects on gene expression in the two cell types (Figure 3). Notably, all SVA eTEs identified in LCL (n=4) were involved in positive correlations (*i.e.* the insertion is associated with increased gene transcript levels) and all were located within intron or proximal to the eGenes (up- or down 10kb) (Figure 3-B). By contrast, in iPSC, SVA eTEs (n=14) were generally associated with negative correlations and mostly distal to genes (X-squared-test, X-squared = 3.8612, *P* < 0.05; Figure 3-A,B). Moreover, 3 out of 4 intronic L1 eTEs were negatively correlated with gene expression in LCL, while 7 out of the 8 L1 eTEs in iPSC display a positive correlation (Figure 3-B). These differences may reflect cell type-specific mechanisms by which the different types of TEs influence adjacent gene expression (see Discussion).

**Figure 3.**
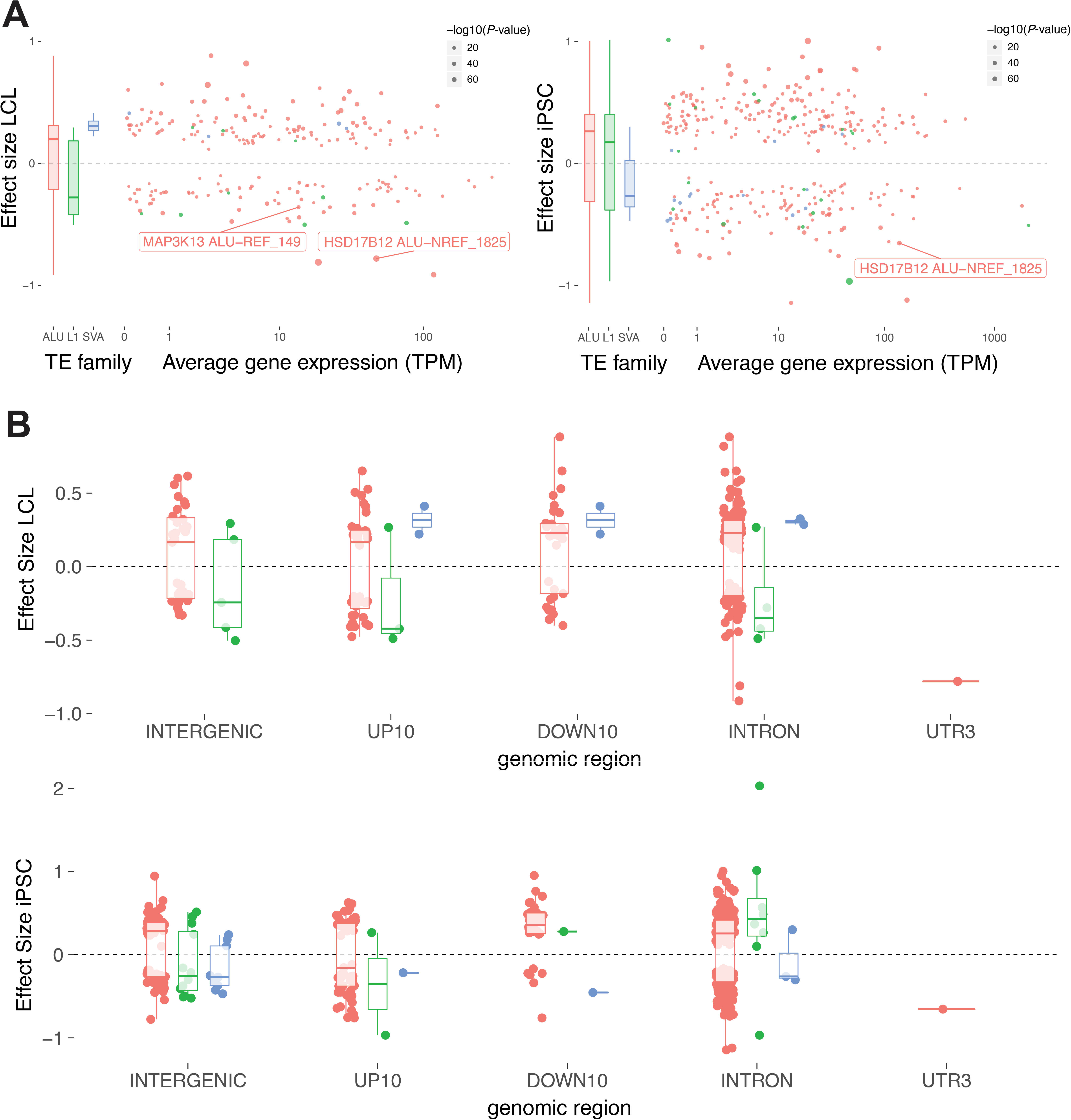
TE-eQTL Effect Sizes. A - Joint distribution of TEI-eQTL effect size (y axis) by eGene average expression (x axis). Negative effect size indicates negative correlation between TE presence and gene expression. Individual loci discussed in the test are boxed. B - direction of the effect sizes for LCL (up) and iPSC (down) relative to RefSeq gene model positions (compartments as defined in legend of Figure 2).

To assess the contribution of eTEs to eQTLs relative to linked SNPs, we performed regional conditional eQTL analysis using SNPs directly located within 1 Mb of each detected eGene (see Methods). One eTE in LCL and six eTEs in LCLs were ranked as the “top-variant” following the procedure implemented in QTLtools (Supplementary Data 1). While the high level of linkage disequilibrium in humans generally obscure the identification of causal variants in QTL and association studies[40], these observations suggest that caution must be used when interpreting the biological impact of TE-eQTLs relative to other linked variants.

### Conservation of TE-eQTL across cell types

Seventy-three (15.3%) of the TE-eQTL were detected in both LCL and iPSC. When applying a Bonferroni correction to the initial *P*-values (threshold = initial *P*-value threshold at 5% FDR / 2), sixty of these shared TE-eQTL remain significant, accounting for 28.4% and 17.7% of the TE-eQTL identified in LCL and iPSC respectively (Figure 4-A). Conversely, cell type-specific TE-eQTL accounted for more than half of the TE-eQTLs identified in each cell type: 52.1% (110/211) in LCL and 63.8% (217/340) in iPSC, after applying Bonferroni correction. The effect size of the 60 shared TE-eQTL were strongly correlated across the two cell types (Figure 4-B, inverse hyperbolic sine transformation, r = 0.87, *P* < 0.01). Nearly all (57/60, 95%) of the shared TE-eQTL also shared the direction of their effect on gene expression (Figure 4-B). We observed that average transcript levels of shared eGenes was also highly correlated across the two cell types (Figure 4-C, Pearson product moment correlation, r = 0.83; P < 0.01). More generally, the average expression levels of eGenes was not significantly different between the two cell types (ANOVA, F = 2.261, *P* = 0.133) nor between shared eGenes and cell type-specific eGenes (Figure 4-D, ANOVA: F = 0.982; *P* = 0.322 [specific *vs* shared], F = 0.176; *P* = 0.675 [interaction ‘cell type’ x ‘specific *vs* shared’]). However, the statistical significance (*P*-value) of shared TE-eQTL was stronger than that of cell type-specific TE-eQTL (Figure 4-E; ANOVA, F = 32.409, *P* < 0.01). In other words, TE-eQTL shared between cell types appear statistically more robust than cell type-specific TE-eQTL, and this distinction cannot be merely explained by differences in basal eGene expression levels.

**Figure 4.**
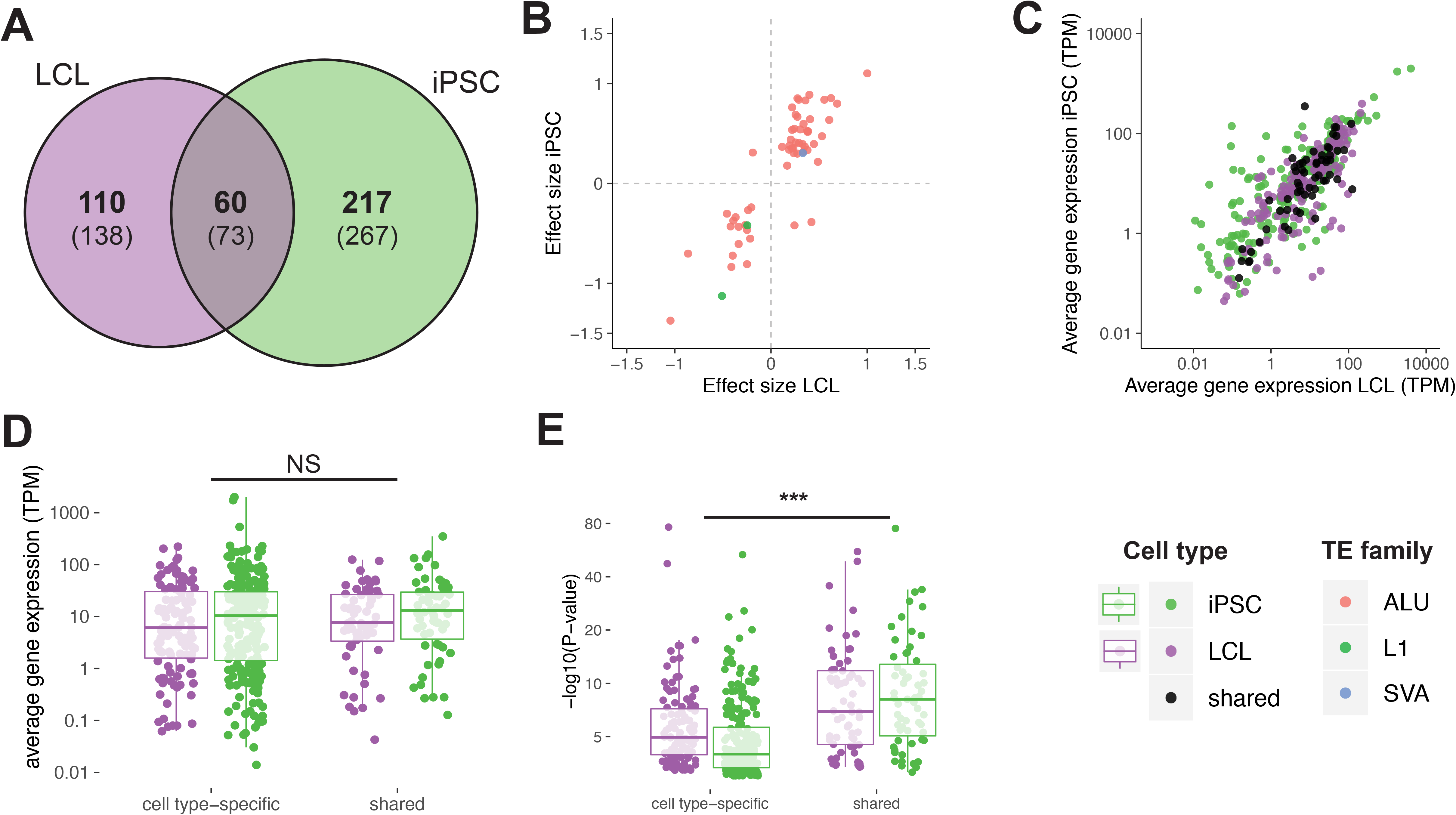
TE-eQTL sharing between cell-type. A - Venn diagram representing the intersection of the total (in parentheses) and significant (in bold) number of eQTL discovered in LCL and iPSC. B - Relationship between the effect size of shared eQTL between LCL (x axis) and iPSC (y axis). C - Relationship between average gene expression in LCL and iPSC for shared and cell type specific eGenes. D - Comparison of average gene expression of cell type-specific and shared TE-eQTL. E - Comparison of P-values distributions between cell type-specific and shared TE-eQTL. Significance: ANOVA, NS: Non significant difference between groups (cell type-specific *vs* shared); ***: *P* < 0.01

Next we examined whether the genes associated with TE-eQTLs (eGenes) were enriched for particular biological functions, states or processes. We used the Gene Set Enrichment Analysis (GSEA) pipeline and datasets (http://software.broadinstitute.org/gsea/msigdb/index.jsp) to analyze eGenes associated with cell-type specific or shared TE-eQTLs using five gene sets precomputed in the GSEA database (H: hallmark gene set, CP: canonical pathways, CP:BIOCARTA, CP:KEGG and BP: GO term Biological Pathways), but restricting each gene set for genes actually expressed in the relevant cell type (Table 1). The results of this analysis revealed that iPSC-specific eGenes were enriched for genes implicated in developmental processes, such as GO_REGULATION_OF_CELL_DIFFERENTIATION (*q*-value = 4.66E-4) and GO_REGULATION_OF_MULTICELLULAR_ORGANISMAL_DEVELOPMENT (*q*-value = 1.74E-3) as well as several housekeeping functions (Supplementary Data 2). However, LCL-specific eGenes did not overlap with immunity-related biological processes as might have been expected if unfixed TEs were to contribute to cell type-specific functions of LCLs. Furthermore, we found that shared TE-eQTL overlapped most significantly with rather broad, housekeeping gene ontologies such as GO_SINGLE_ORGANISM_BIOSYNTHETIC_PROCESS (*q*-value = 4.85E-3), GO_CELLULAR_AMINO_ACID_METABOLIC_PROCESS (*q*-value = 3.71E-2) as well as the stress response (GO_CELLULAR_RESPONSE_TO_STRESS, *q-*value = 3.71E-2) (Supplementary Data 2).

In summary, we found that a substantial fraction of TE-eQTL overlap between the two cell types considered, and their statistical significance tend to be stronger than that of cell type-specific TE-eQTL. Additionally, we found no striking association of cell type-specific TE-eQTL with genes involved in highly specific pathways or ontologies, though we note a modest enrichment of iPSC-specific TE-eQTL with developmental genes.

### Chromatin accessibility TE-QTL

To shed light into the mechanisms by which unfixed TEs may regulate gene expression at the chromatin level, we leveraged ATAC-seq data available for a subset of 86 LCL from the GBR population[41] to perform a chromatin-accessibility QTL (caQTL) analysis. Using the TE genotypes predicted for these cell lines, the analysis yielded a total of 656 significant TE-caQTL (FDR 5%, with 10,000 permutations of the ATAC peaks quantification values), involving 431 distinct caTEs (Figure 5-A). Nearly 90% (585/656) of the predicted interactions between caTE and associated ATAC peaks occur within 250 kb (Figure 5-B), a distance compatible with a direct effect of the TE on accessibility of the regulatory DNA through chromatin looping or spreading [42, 43]. We also observe that the distance between caTE and associated ATAC peaks follows a distribution that is slightly broader (F-test, F = 0.85623, *P* = 0.0472) than the one obtained with a matching number of SNP-caQTL, but is indistinguishable from the distance distribution between eGenes and eTEs mapped in the LCL (F-test, F = 0.91358, *P* = 0.4061).

**Figure 5.**
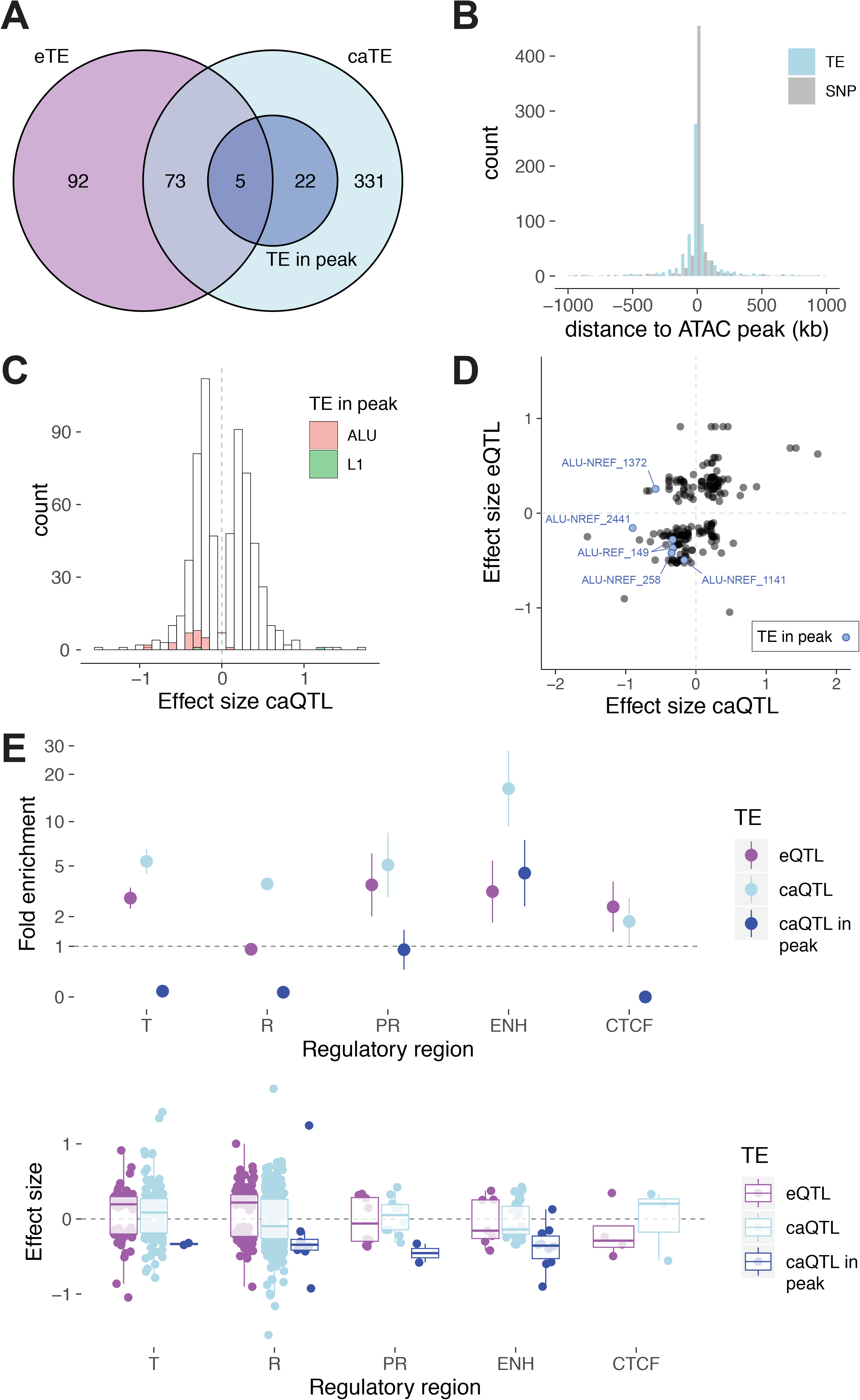
Chromatin accessibility QTL in LCL (TE-caQTL). A - Venn diagram representing the overlap between eTE, caTE and caTE directly mapping in the associated ATAC peak. B - Distribution of the TE-caQTL size effect. Families of caTE inserted directly within the associated ATAC peak are colored according to the legend. C - Distribution of the distance between caTE and their associated ATAC peaks. D - Relationship between eQTL and caQTL effect sizes for 78 TE involved in the two QTL. Blue dots are caTE inserted in the associated ATAC peak. E - Enrichments of eTE and caTE in different regulatory regions according to the ChroMM+Segway integrated track generated for one of the LCL (GM12878). T = transcribed region; R = repressed region; PR = Promoter (includes TSS); ENH = Enhancer; CTCF = CTCF binding site.

Interestingly, seventy-eight caTE-QTL (18%) were also detected as involved in at least one eTE-QTL in the LCL (Figure 5-A), a much greater overlap than expected by chance if the two were independent (X-squared test, X-squared = 131.61, *P*-value < 2.2e-16). Such overlap raises the possibility that some of these eTEs affect gene expression by modulating chromatin accessibility of nearby *cis*-regulatory elements. Consistent with this model, the 78 TEs involved in both eQTL and caQTL display a faint but significant correlation of their effects on each type of QTL (Figure 5-D, Pearson’s product moment correlation test, r = 0.47, *P* < 0.01).

A particularly interesting subset of TE-caQTL are those where the TE directly overlaps with its associated ATAC peak (dark blue, Figure 5-A), which likely implies that the TE insertion itself is responsible for the modulation of chromatin accessibility. Strikingly, 25 out of 27 such caTEs mapping within ATAC peaks were associated with reduced peak size (*i.e.* negative effect size, Figure 5-C,D). These data suggest that the insertion of a TE within a *cis*-regulatory element generally lowers its accessibility, which in turn could lead to repressive effect on adjacent gene transcription. Indeed, out of the five caTEs within peaks which were also associated with eQTLs, four were associated with reduced eGene expression (Figure 5-D). These data support a model whereby the insertion of TEs within *cis*-regulatory elements is generally disruptive and often leads to reduced transcription of their target genes.

An evocative example is an AluYb8 element inserted within the third intron of the *MAP3K13* gene (Figure 6). The presence of the TE correlates with reduced chromatin accessibility at three ATAC peaks surrounding the gene, including one predicted as a transcribed enhancer in LCL directly overlapping with the TE, and the same TE insertion was also associated with reduced *MAP3K13* expression in our TE-eQTL analysis (Figure 6). Because *MAP3K13* is known to be a positive regulator of the proto-oncogenic transcription factor *c-Myc*[44], it is tempting to speculate that the AluYb8-containing allele might confer an anti-tumorigenic effect by attenuating *MAP3K13* expression (see Discussion).

**Figure 6.**
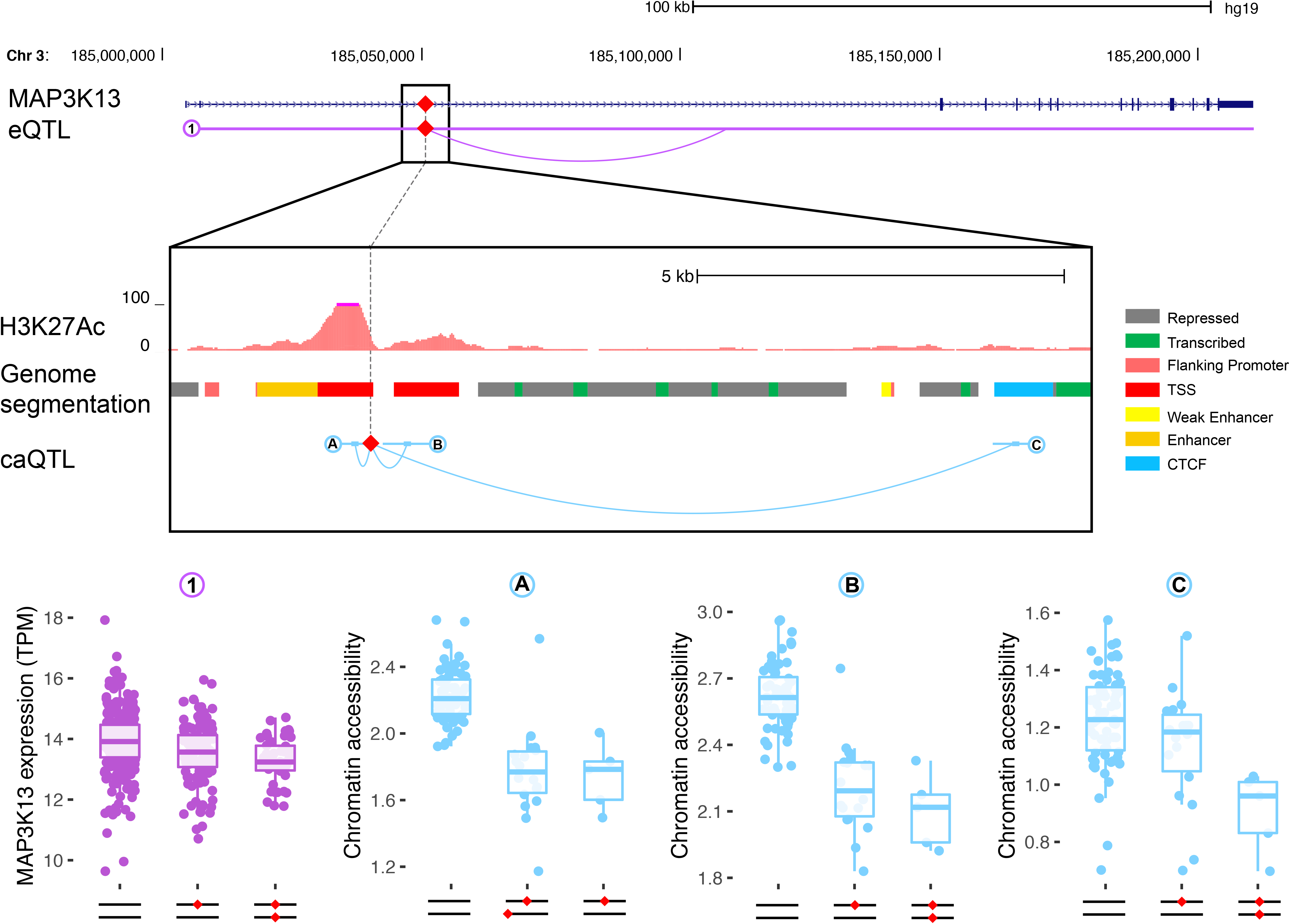
Unfixed AluYb8 insertion correlates with chromatin accessibility and gene expression of *MAP3K13*. The insertion of an AluYb8 (red diamond) in a putative enhancer within the second intron of the *MAP3K13* gene is both associated with reduction in *MAP3K13* mRNA levels (1) and reduced chromatin accessibility at three ATAC peaks (A, B and C). The figure reproduces the H3K27Ac track from ENCODE for the LCL GM12878; Genome segmentation according to figure legend from the combined ChromHMM+Segway ENCODE track for GM12878.

### Post-transcriptional effects

To identify plausible cases of post-transcriptional effects of unfixed TEs on gene expression, we focused on 9 TEs included in our original dataset located within the 3’-UTR of genes. We found that only one was a significant TE-eQTL: an AluYa5 element (ALU-NREF_1825) inserted within the 3’-UTR of the major transcript for *HSD17B12.* This transcript encodes for a hydroxysteroid 17-beta dehydrogenase involved in long chain fatty-acids elongation and its expression level has been associated with cancer prognosis in human and fertility in mice[45–47] (Figure 7). This was one of the strongest TE-eQTL mapped in both LCL and iPSC (Figure 3-A). In both cell types, mRNA levels of *HSD17B12* were negatively correlated with the presence of the Alu insertion (Figure 7-A). Regional eQTL analysis including TEs, SNPs and indels (see methods) revealed a strong linkage block, with markers reaching the highest *P*-values within the 3’-UTR of *HSD17B12* (Figure 7-B). Conditional analysis of these significant markers (including SNPs, indels and TEs) ranked ALU-NREF_1825 respectively 1/340 (top variant) and 74/346 in iPSC and LCL, respectively. Together these data strongly implicate this Alu insertion in modulating the expression of *HSD17B12*.

**Figure 7.**
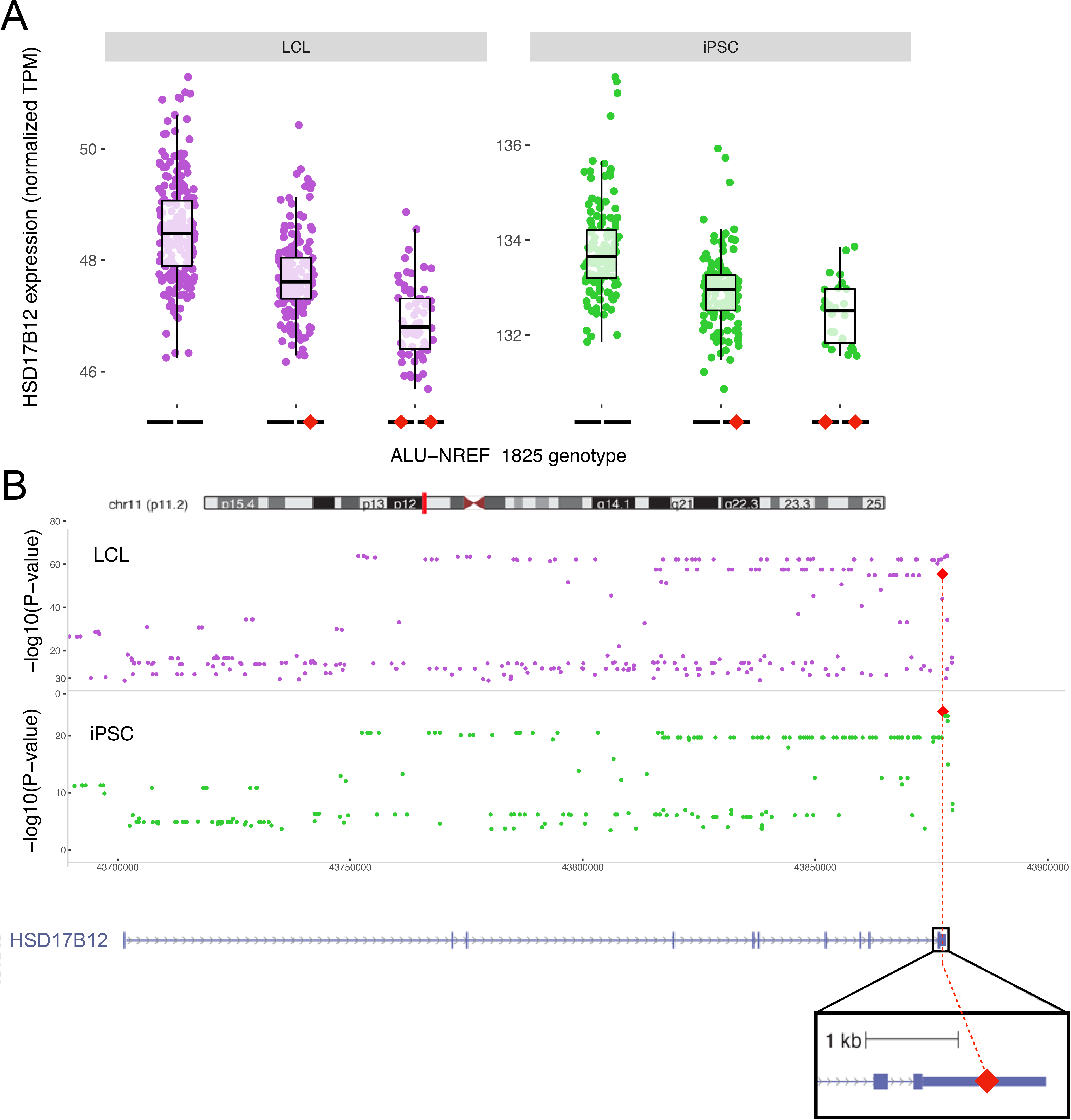
Unfixed AluYa5 insertion within the 3’-UTR of *HSD17B12* is associated with transcript downregulation. A - Boxplots comparing the expression of the gene HSD17B12 (RNA-seq) according to the TE genotype (the TE insertion is represented by a red diamond) in LCL (left) and iPSC (right). B - Regional analysis of unfixed TE and other variants (SNP) associated with the expression levels of HSD17B12. Top: LCL, bottom: iPSC (Alu is top variant).

To investigate whether the Alu insertion within the 3’ UTR could affect protein expression and whether this effect is determined by the Alu sequence, we utilized a luciferase reporter assay designed to compare the effects of three different 3’-UTRs cloned downstream of the luciferase coding sequence (Figure 8): (i) a 3’-UTR sequence derived from an allele lacking the insertion (from individual NA11830); (ii) a 3’-UTR sequence derived from an allele containing the AluYa5 insertion (from individual NA12760); (iii) a 3’-UTR sequence derived from the same Alu-containing allele but with a randomly scrambled sequence of the Alu (see Supplementary Data 3 and Methods). Reporter plasmids were transfected by electroporation into the LCL GM12831 (NA12831) in order to perform the assays in a cellular environment comparable to that of the eQTL analysis. The results show that attaching any of the three 3’-UTR to the luciferase coding sequence leads to a significant decrease in protein expression compared to a construct without any 3’ UTR, but downregulation was significantly greater for the two Alu-containing constructs, regardless of whether the Alu sequence was scrambled or not (Figure 8). These results recapitulate the eQTL data and suggest that the presence of the AluYa5 insertion downregulates gene expression most likely at the post-transcriptional level, but this effect is apparently independent of the Alu sequence.

**Figure 8.**
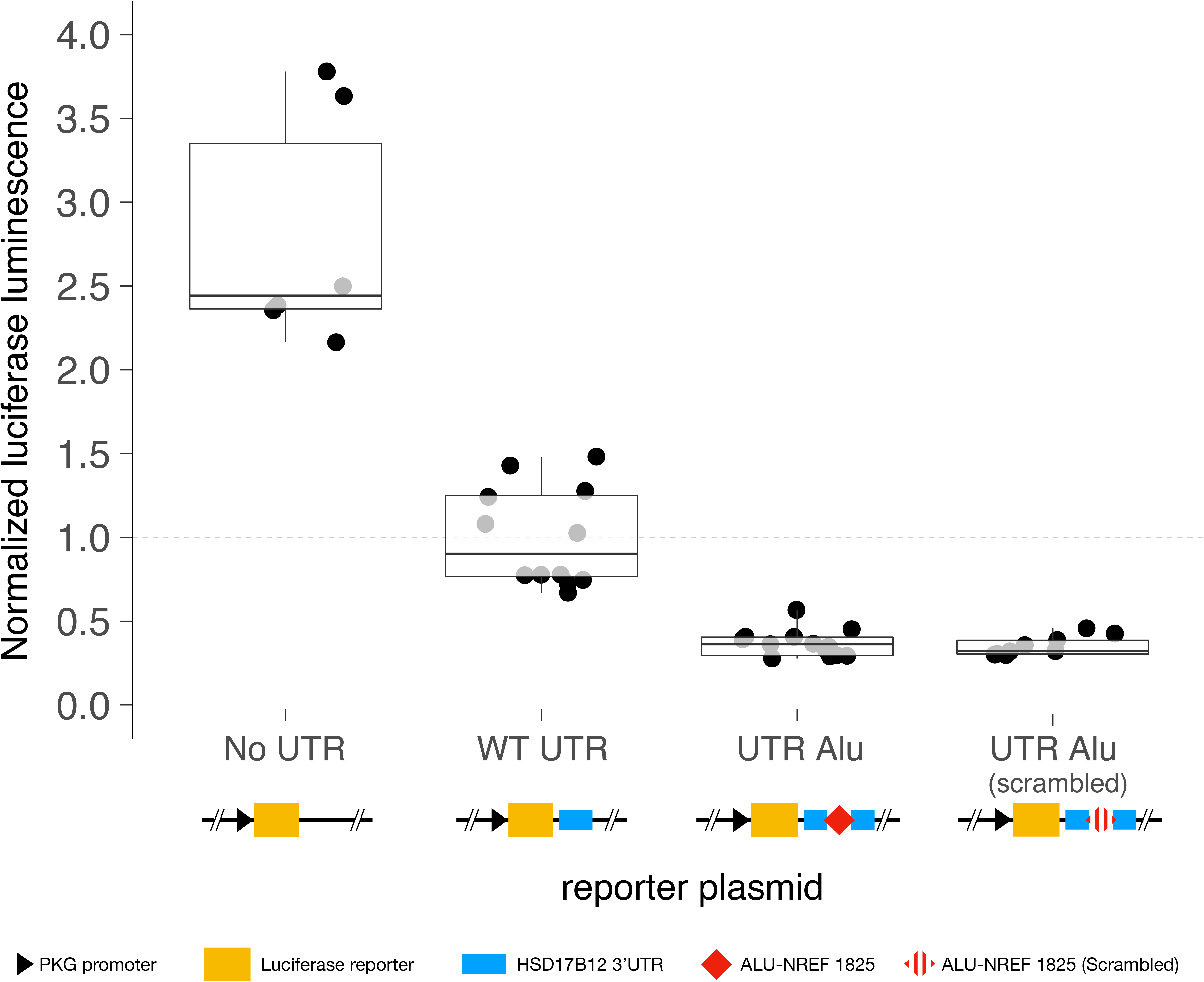
Alu-bearing 3’-UTR haplotype of *HSD17B12* show reduced reporter protein expression levels. Relative luciferase luminescence of transfected LCL (GM11831) with different 3’UTR construct: “No UTR” = empty plasmid; “WT UTR” = UTR without Alu insertion (from NA11830); “UTR Alu” = UTR with Alu insertion (from NA12760); “UTR Alu (scrambled)” = UTR with a scrambled sequence of the AluYa5 present in NA12760.

## Discussion

Unfixed TE insertions represent an important class of structural variants between human genomes, but their impact on gene regulation remains poorly characterized [16–19]. To our knowledge, this study is only the second to broadly assess the regulatory potential of unfixed TEs on human gene expression [18] and the first to consider their effects across multiple cell types. We also present the first TE-chromatin accessibility QTL (TE-caQTL) analysis, which sheds light on the impact of recent TE insertions on chromatin state at or near their insertion sites and enables a more direct evaluation of their contribution to *cis*-regulatory variation.

By leveraging newly predicted genotypes for 2,806 TE insertions, we were able to identify 211 *cis*-TE-eQTL across 444 LCL and 314 *cis*-TE-eQTL across 294 iPSC. A previous analysis of the same LCL dataset using a similar analytical framework reported 53 *cis*-TE-eQTL (outside the HLA and 17q21.31 loci) [18], including 20 loci that our analysis also identified. The difference in the outcomes of the two studies, and notably the considerably larger set of TE-eQTLs captured by our approach, can be attributed to several important methodological differences. First, we used an improved genotyping pipeline, TypeTE [31], allowing us to consider 860 unfixed TEs present in the reference genome assembly (GRCh37/hg19)[12], which were not included in the dataset used in previous studies[17,18,32] likely due to the unreliability of the original genotypes provided by the 1000 Genome Project ([12, 31]). A second major difference is that we re-processed the raw RNA-seq reads originally produced by Lappalainen et al. [35] to match the more recent quantification procedures adopted by the HipSci consortium (Helena Kilpinen, personal communication, www.hipsci.org). Additionally, we employed a recently developed QTL mapping toolkit, QTLtools, which relies on a more robust statistical framework than previous approaches [36]. Together these improvements likely increased the power of our analysis and enhanced our ability to map TE-eQTL with these datasets.

Consequently, we were able to explore finely the regulatory potential of unfixed TEs across two different human cell types. In both LCL and iPSC, we found that most predicted *cis*-regulatory interactions take place within 250 kb of the gene boundaries, a value consistent with eQTLs previously mapped with SNPs [21, 33], as well as our own eQTL analysis using SNPs (Figure 2-A,B). We also noticed a trend for *P*-values to weaken as the distance between eTE and eGene increases (Figure S.4). These observations are consistent with the current understanding of how distal *cis*-regulatory elements interact with genes in the human genome[48, 49], within topological associated domains of a median length of ∼185 kb [43]. Repeating the analysis with increasing subsamples (Figure S.2) suggests that many more TE-eQTL remain to be discovered in the human population. This finding is consistent with previous large-scale SNP-eQTL studies [21, 25].

Our study provides a first assessment of whether a TE-eQTL discovered in a given cell type may be detected in another. We found that 28% of the TE-eQTL identified in LCL were replicated in iPSC, representing nearly 18% of the significant TE-Gene associations in the latter cell type. The level of statistical significance, as well as the size and direction of the effect on gene expression, were highly correlated across the two cell types, suggesting that their functional impact may be broadly conserved. These findings suggest that many TEs could influence gene expression across multiple cell types and a potentially wide range of tissues [21, 50]. Likewise, SNP-eQTL analyses across 44 human tissues by the GTEx consortium suggest that SNPs with *cis*-regulatory effects fall within one of two broad categories: those with shared effects across tissues and those specific to a single or a few similar tissues [21]. Our results suggest that this dichotomy may also apply to TE insertions, which implies that the phenotypic effect of a TE insertion in a given tissue could be inferred from data obtained from another tissue.

While the eGenes detected in our analysis may not necessarily serve cell type-specific functions (Supplementary Data 2), we observe interesting trends in the *cis*-regulatory effects associated with different types of TEs across the two cell types. Notably, SVA-eTEs mapped in LCL were all correlated with increased gene expression, while SVA-eTEs mapped in iPSC were mostly associated with decreased gene expression (Figure 3). Intronic L1-eTE showed the opposite trend: they tend to associate negatively in LCL but positively in iPSC. These trends may reflect intrinsic differences in the *cis*-regulatory potential of these TE families. Indeed, while SVA is mobilized by the L1 machinery, their expression pattern are only partially overlapping and each family appears to exhibit unique cell type-specific regulatory activities [51–54]. Taken together, our findings indicate that the *cis*-regulatory effects of unfixed TEs may often manifest in multiple cell types, but further investigation is needed to characterize possible cell type-specific interactions.

One of the novelties of our study is to incorporate functional genomics data to investigate further the regulatory potential of unfixed TEs. To our knowledge, we report the first TE-chromatin accessibility QTL (TE-caQTL) analysis. We used ATAC-seq data generated previously for 86 of the LCL [41] to map 431 caTEs associated with variation at 656 ATAC peaks. Strikingly, more than one third of the TEs involved in eQTL were also mapped as caQTL, which brings support to the hypothesis that a substantial fraction of TE insertions modulate adjacent gene expression through distal chromatin effects, as previously proposed for SNP-eQTL in LCL [55] and iPSC [50]. Furthermore, we found that TE associated with chromatin accessibility peaks were ∼4 times more enriched within regions annotated as enhancers than TE-eQTL, which is consistent with the idea that some of the TE insertions detected in the TE-caQTL analysis could directly modify or act as *cis*-regulatory element [56, 57]. As a support for this hypothesis, we found that 27 of the caTEs were directly located within their associated ATAC peak and all but two of these insertions correlated with reduced chromatin accessibility. The results of our TE-caQTL analysis indicate that TE insertions within *cis*-regulatory elements, such as promoter or enhancers, tend to have disruptive effect on the function of these elements.

It is well documented that TEs can affect post-transcriptional gene regulation through many mechanisms, including effects on mRNA splicing, stability or translation[2,4,19,58,59]. While there are many known examples of fixed human TEs with such effects, cases involving unfixed elements have been scarcely described outside of disease-causing insertions [8,9,60]. We confirmed experimentally that the inclusion of an Alu in the 3’-UTR of the gene *HSD17B12* (previously reported by Wang et al. [18]) reduces the protein level of a luciferase reporter in LCL. However, this effect appears to be sequence-independent, since a reduction in luciferase expression was observed even when the Alu sequence was scrambled. These results seem to rule out some of the known mechanisms by which (fixed) Alu located in 3’-UTRs affects transcript stability, such as miRNA binding [58] or Staufen-mediated decay [61]. It is possible that that the effect merely reflects the elongation of the 3’-UTR caused by the insertion. Indeed, it is known that the RNA helicase Upf1 can sense 3’-UTR size and promote nonsense mediated decay of abnormally long 3’-UTRs [62]. Also, because we cloned the “presence” and “absence” 3’-UTR haplotypes from two different LCLs, we cannot rule out that another polymorphism ‘hitchhiking’ with the Alu insertion, is causing the effect. At the very least, our results indicate that the Alu insertion can act as a reliable marker of HSD17B12 expression. Because the level of HSD17B12 enzyme has been positively correlated to the severity of epithelial ovarian cancer in humans [45], this is a case worth further investigation.

The example of *HSD17B12* and the newly reported case of *MAP3K13* (where an Alu insertion could be mapped both as expression and chromatin accessibility QTL in LCL) as well as the many other TE-QTLs identified in this study underscore a plausible contribution of TE insertions commonly segregating in the population in human trait variation, including disease susceptibility [16,19,63]. These findings confirm the potential of unfixed TEs to make a non-trivial contribution to human gene expression [5–7]. Our study provides evidence for the persistence of their (putative) *cis*-regulatory effects across cell types, but shows that some elements have the potential to regulate tissue-specific functions. Moreover, we present the first map of unfixed TEs that correlate with changes in chromatin accessibility in *cis*, uncovering the importance of this mechanism while its fine details remain to be investigated. A logical extension of this study would be to leverage data produced for a broader range of human tissues, such as those represented in the GTEx initiative [21] to analyze more comprehensively the tissue-specificity of the TE-eQTL identified herein. Complementary genomic assays, such as those measuring DNA methylation levels or nascent RNA transcription, could provide further insight into the mechanisms by which polymorphic TE insertions shape gene expression. Ultimately, the causality of TE insertion variants would need to be tested experimentally through CRISPR-cas and other manipulative genomic technologies [5]. The data presented here offer a valuable foundation for future studies aimed at illuminating the contribution of TEs to human phenotypic variation.

## Methods

Unless otherwise stated, all statistical analyses were performed using R version 3.5.1 (R development core team, 2018).

### TE genotypes

The genomic locations of unfixed Alu, LINE1 (L1) and SVA element were extracted from publicly available datasets. TE insertions in LCL were gathered from the previous analysis of 445 cell lines derived from healthy donors of 5 populations (CEU, FIN, TSI, GBR and YRI) by the 1000 Genome Project[12]. TE insertions were originally discovered and genotyped in this dataset using MELT v1 (non-reference insertions) and a collection of structural variant tools[12]. In some cases, these calls were re-genotyped using TypeTE (Goubert et al. 2019, BiorXiv) in order to improve their accuracy. TypeTE was used for L1 and SVA insertions present in the reference genome, as well as for both reference and non-reference Alu insertions. In addition, unfixed TE insertions were searched and genotyped *de-novo* in 326 induced human pluripotent stem cells (iPSC), derived from 205 healthy donors as follow. First, whole genome sequencing data for each cell line was recovered in bam format from the HipSci website (HipSci.org) and unfixed TE insertions were called using MELT version 2.1.4[15] using split- and discordant paired-end read information. Since genotype accuracy has substantially improved for non-reference Alu between MELT1 and MELT2 (Goubert et al, 2019 BiorXiv), we only re-genotyped L1, SVA as well as reference Alu insertions. To match insertions found in both datasets, we intersected the breakpoint positions of the polymorphic insertions discovered in LCL and iPSC using bedtools (version 2.28.0 [64]). Two insertions of the same TE type (Alu, L1 or SVA) separated by up to 30 bp were considered identical by descent. After genotyping, only shared loci with a minimum insertion frequency of 5% in each dataset were kept for eQTL mapping.

### RNA-seq data processing

Steady-state RNA levels for LCL and iPSC were recovered and normalized as follow. We collected reads from the RNA-seq experiment carried-out by Lappalainen et al.[24] in 445 LCL from the Geuvadis repository (https://www.ebi.ac.uk/Tools/geuvadis-das/). Raw reads were quality checked and trimmed using UrQt (version 1.0.18, [65]); we used a -t quality threshold of 10 and kept the other default parameters. Transcript levels were then quantified with kallisto (0.46.0, [34]) using the reference transcriptome (cDNA) GRCh37.75 from Ensembl. This reference transcriptome and quantification method were used to match the data of 326 iPSC (Kilpinen, personal communication, HipSci.org). Sample quantifications in transcript per million (TPM) were then grouped by cell-type and normalized. TMM (trimmed mean of M-values) normalization was performed to make transcript level comparable across samples using the script abundance_estimates_to_matrix.pl available with the Trinity distribution version 2.8.4[66]. Transcript quantifications were summed by gene and those expressed in less than 50% of the samples were then discarded. Then, PCA using normalized TPM were carried out to identify outlier samples. Samples with values exceeding 3 times the standard deviation on each of the first 2 principal components were removed. After filtering, 444 and 294 samples were kept respectively in LCL and iPSC.

### TE-eQTL mapping

TE-eQTL were mapped independently in LCL and iPSC datasets using QTLtools v1.1 [36]. After ensuring than transcript expression was not structured by population in the LCL dataset, we used QTLtools to perform a new PCA analysis on the final expression matrices. For each cell-type, the values of the three first axes were added as covariates to the model, as well as the sex and population of origin for LCL and sex, ethnicity and age for iPSC. *Cis*-eQTLs were searched within a 1 Mb window around each transcript using QTLtool *cis*. Significance was evaluated by running 10,000 permutation of the gene expression matrices and multiple testing was addressed by applying 5% FDR correction, as recommended in the QTLtools manual. The top eTEs (most significant TE insertion significantly associated to a gene expression level) reported by QTLtools were kept for further analysis. Our ability to detect TE-eQTL was evaluated by resampling an increasing number individuals selected at random (10, 25, 50, 100 and 200) in the TE genotype matrices and re-running the QTLtools *cis* procedure. Enrichments of TEs in specific gene regions (intergenic, 10 kb upstream, 10 kb downstream, intron, exon, 5’UTR or 3’UTR) were calculated for all TEs within the 1 Mb window around a given gene and eTEs reported by QTLtools. Enrichments were calculated by sampling a matching number of 1 bp random genomic interval (TE breakpoints) in the reference genome using bedtools for 1,000 times. The ratio observed TE / random breakpoints in a given region was then calculated for each replicate to calculate a fold enrichment.

### TE-eQTL sharing between cell types

Sharing of TE-eQTLs between cell-type was considered significant if two identical gene-TE associations had a *P*-value below or equal to half the initial FDR threshold (Bonferroni correction). Sharing was also quantified in re-sampled eQTL analyses (increasing sample numbers) using the same criteria.

### Conditional analysis of TE- and SNP-eQTLs

The regulatory potential of polymorphic TEs was compared to SNPs by performing conditional eQTL analysis [36]. The SNP dataset used for this analysis was recovered from the 1000 Genome Project phase 3 release for LCL [67], while for iPSC we used individual vcfs files available from the HipSci consortium (these include imputed 1000 Genomes genotypes). To relieve the computational burden of mapping eQTL for all SNP, we extracted only markers present in the 1-Mb windows where a TE-eQTL has been previously detected. Combining TE and SNP genotypes, eQTL were searched in each cell type using QTLtools *cis*. A first pass was performed using 10,000 permutations with a FDR threshold of 5%, then, a conditional analysis for each gene was performed in order to rank the e-variants (variant, either TE or SNP, associated with a gene expression level), using the “--mapping” option of QTLtools *cis*.

### TE-caQTL mapping

To investigate potential cis-effects of TE insertions on chromatin accessibility, we collected ATAC-seq data for 85 GBR individuals included in our LCL dataset[41]. Normalized ATAC peak levels were used as response variable to search for TE-caQTL with QTLtools, using the genotypes of unfixed TEs for these LCL. As for eQTL mapping, we used population and sex of each cell line as covariate and report the best significant TE per normalized ATAC peak as provided by QTLtools in a cis-window of 1 Mb. The resulting caTEs (significant TE in caQTL) were then searched for enrichment in regulatory regions by intersecting their genomic location with the Segway/ChromHMM combined regulatory track from ENCODE generated for the LCL GM12878.

### Luciferase reporter assays

#### UTR amplification

We evaluated the regulatory potential of a candidate AluYa5 insertion within the 3’UTR of the gene HSD17B12 using dual luciferase reporter assay. First, three LCL corresponding to one homozygote for the insertion (GM12760), one homozygote for the absence (GM11830) and one heterozygote (GM12831) were cultured in RMPI (Gibco) supplemented with 15% FBS (Gibco) at 37C and 5% CO2. Cells were passagedmost significant and refreshed with new media every 3 to 4 days, before reaching ∼1 million cell per milliliter of culture. 3’ UTR sequences with and without Alu insertion were amplified from homozygote individuals by PCR as follow. DNA was extracted using Qiagen© DNeasy Blood and Tissue kit following manufacturer instructions. For PCR amplification, no more than 1 ug of DNA template was used per sample in a total volume of 25 µL. 5µL of Q5 High Fidelity master mix (2X) was added with 0.3 µL of each primer (F 5’- **AAACGAGCTCGCTAG**TCAAACCTGCCTTCTTGGA- 3’ and R 5’- **CGACTCTAGACTCGA**CTGTCCAGGTCATTGTGGTG -3’). Primer includes 15 bp homology with the reporter plasmid, as needed for inFusion-HD cloning (Takara Bio Inc). The mixture was amplified after 30 seconds initial denaturation for 25 cycles (10 seconds denaturation at 98C, 30 seconds annealing at 60C and 20 seconds elongation at 72C) followed by 2 minutes of final elongation. Additionally, a construct similar in nucleotide composition to the UTR amplified from GM12760 was generated with a scrambled Alu sequence instead of the original transposable element (supplementary data 3). These three sequences were respectively named “WT UTR” (GM12830, no Alu in UTR), “Alu UTR” (GM12760, Alu in UTR) and “Alu UTR scrambled” (UTR identical to GM12760 but Alu sequence scrambled). In order to assess successful amplification of the UTRs, the band corresponding to the expected PCR product (1,551 bp for “WT UTR” and 1,825 bp for “Alu”) were subject to Sanger sequencing using the F- and R-m13 primers flanking the insert as well as two internal primers (H3Pint1-F: 5’-CAGACACACTGCAATTTACAAAGA-3’ and H3Pint1-R: 5’-ACGGCCTTAATTTCAATCACCA-5’) to fill the gap. PCR products with the expected sequences were then kept for In-Fusion cloning into the reporter plasmid.

#### In-Fusion Cloning into reporter plasmid

The artificially generated “scrambled” sequence was also amplified using the same PCR condition as the natural UTRs to add the 15bp flanking sequences matching the cloning site of the receiving vector. PCR products were then cloned into a pmirGlo dual-luciferase miRNA target expression vector (Promega). This plasmid contains a multiple cloning site downstream of the luciferase gene, terminated by a polyadenylation signal. As a transfection control, the plasmid also contained a renilla reporter gene whose expression should not be affected by the sequence cloned downstream the luciferase. Cloning was done using infusion-HD cloning kit (Takara Bio inc) following provider documentation. Plasmids were transformed into competent alpha-5 E. coli (New England Biolabs) and cultured on LB-Agar plates with 1% Ampicillin overnight at 37C. Next, 10 clones per condition were extracted using mini-prep kit (Qiagen) and successful insertion of the constructs were assessed by preforming double digest of the plasmid by BamHI and EcoRI. Clones with the expected product size (“WT UTR”: 5,681 and 3,188 bp; “UTR Alu” and “UTR Alu scrambled”: 5,955 and 3,188 bp) were then selected for the luciferase reporter assay and amplified by maxi prep (Qiagen) upon transfection.

#### Luciferase assay

“WT UTR”, “UTR Alu”, “UTR Alu scrambled” were transfected into the LCL GM11831 for reporter assay, as well as a pGFP reporter plasmid to evaluate transfection efficiency. For each condition and each replicate, 50 million cells were transfected with 125 ug/µL of plasmid by electroporation using Neon transfection system (Life Technology). Reaction took place in 100µL tips, applying 3 pulses of 1200 V for 20 ms each. Transfection efficiency were ∼30%, and luciferase signal was above background (pGFP cells) by two orders of magnitude for the conditions tested. Reporter assay was performed using Dual-Glo Luciferase assay system (Promega) following manufacturer instructions. For each experiment, the average blank (pGFP) values were subtracted from each luminescence signal. Additionally, luciferase signal was normalized for each replicate by the renilla luminescence, and the ratio luciferase/renilla were eventually normalized by the average ratio of the “WT UTR” construct.

## Acknowledgments

We would like to warmly thanks Mitchell Lokey and Kathleen Gordon for their sustained help during this study. We are grateful to Kathleen Burns and Lindsay Payer for sharing preliminary data and insights to this project. We also thanks Andrew Grimson, Ciáran Daly and Jessica West for their help and providing reagent for the luciferase assay.

