## Supplementary Figures for "Contribution of unfixed transposable element insertions to human regulatory variation"

**Figure S. 1. QQ-plot of TE-eQTL P-values.**

**Figure S.2. TE-eQTLs found as a function of sample size for LCL and iPSC**

**Figure S.3. Effect size as function of the distance between eTE and eGene**

**Figure S.4. TE-eQTL -Log10(*P*-value) as function of the distance between eTE and eGene**

**Figure S.5. TE-eQTL Effect size as function of -Log10(*P*-value)**

**Figure S.6. PCA of 444 LCLs based on normalized gene expression**

Supplementary Data:

Sup. Data 1. TE-QTL results

Sup. Data 2. Gene set enrichments

Sup. Data 3. UTR scrambled sequence (.fasta)

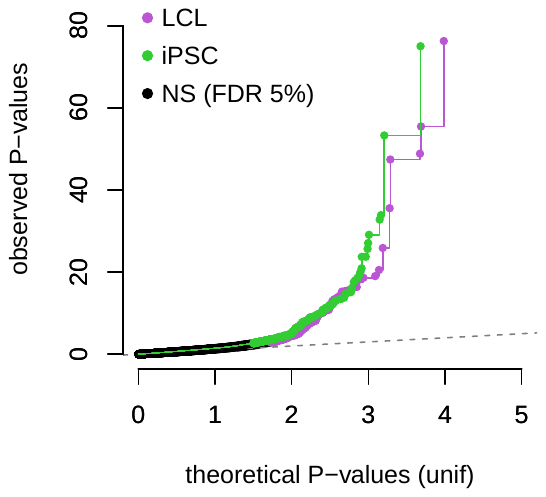

**Figure S. 1. QQ-plot of TE-eQTL P-values.**

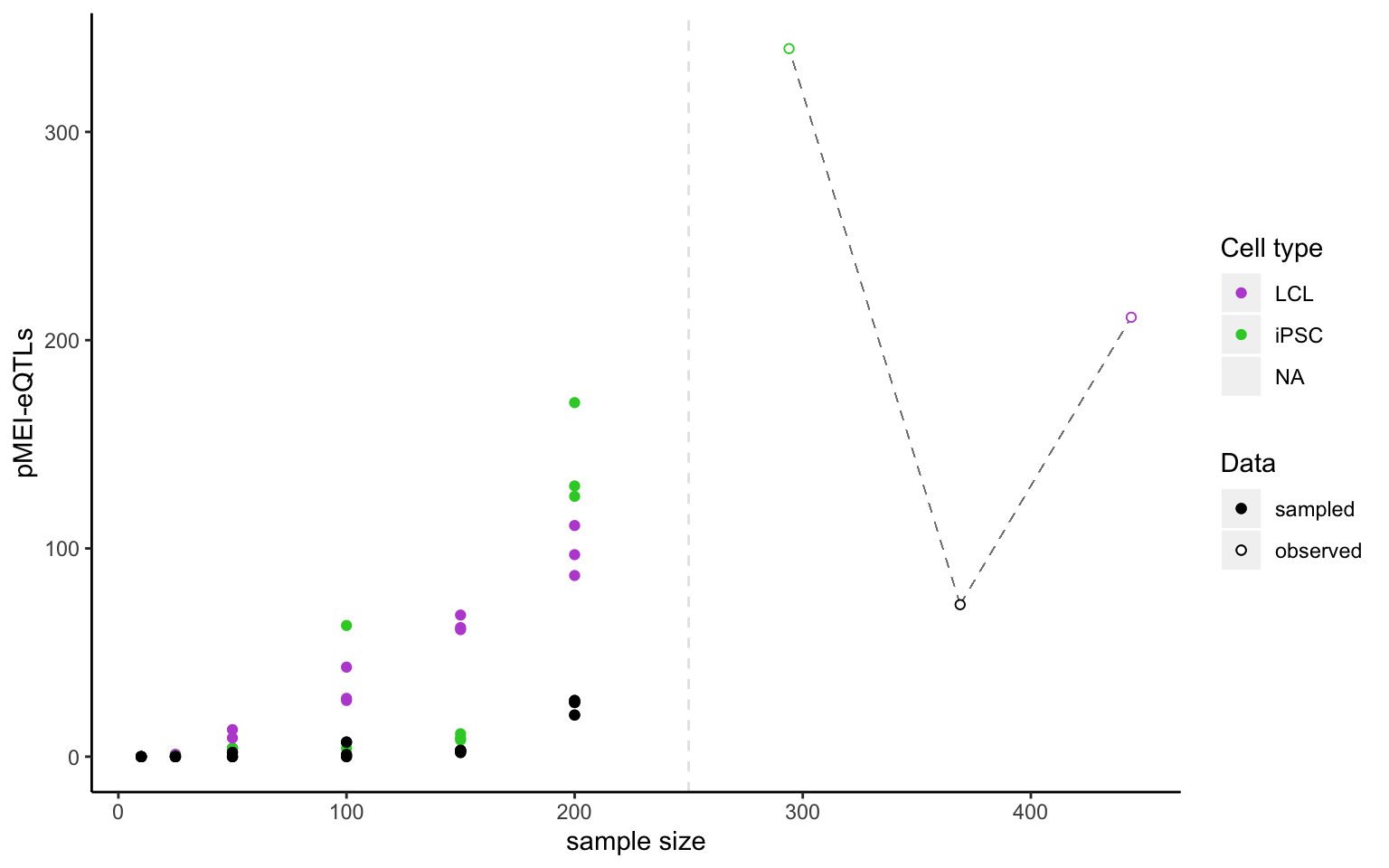

**Figure S.2. TE-eQTLs found as a function of sample size for LCL and iPSC.** Individuals of each dataset are randomly sampled for each analysis. Shared TE-eQTL found between LCL and iPSC are also reported. Resampled value are presented on the left and observed value on the right of the dashed line

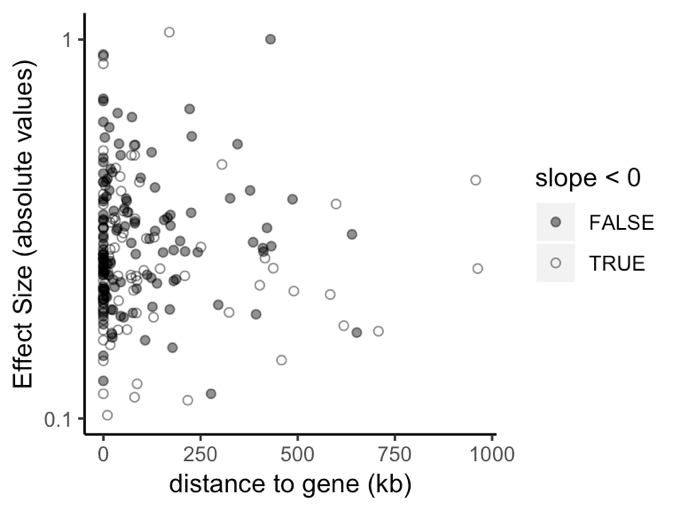

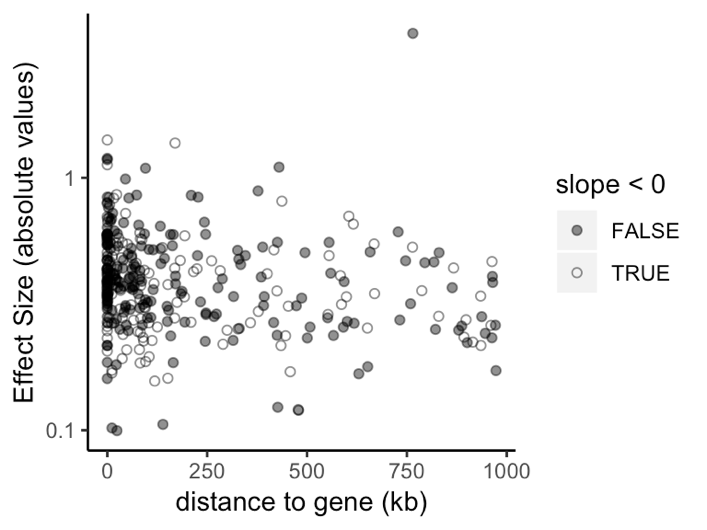

**Figure S.3. Effect size as function of the distance between eTE and eGene** in LCL (left) and iPSC (right)

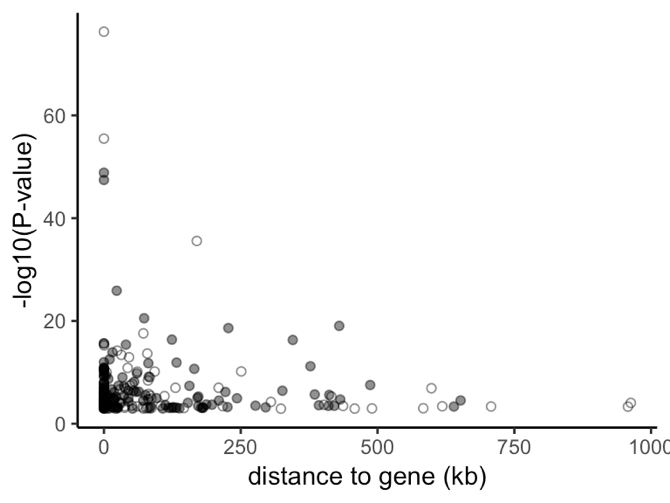

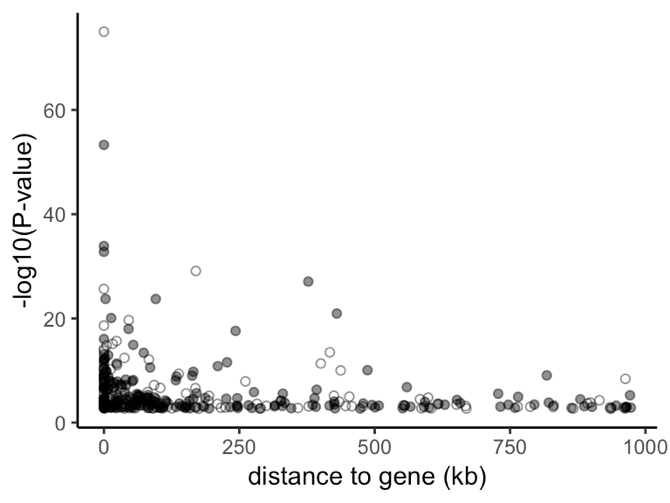

**Figure S.4. TE-eQTL -Log10(*P*-value) as function of the distance between eTE and eGene** for LCL (left) and iPSC (right)

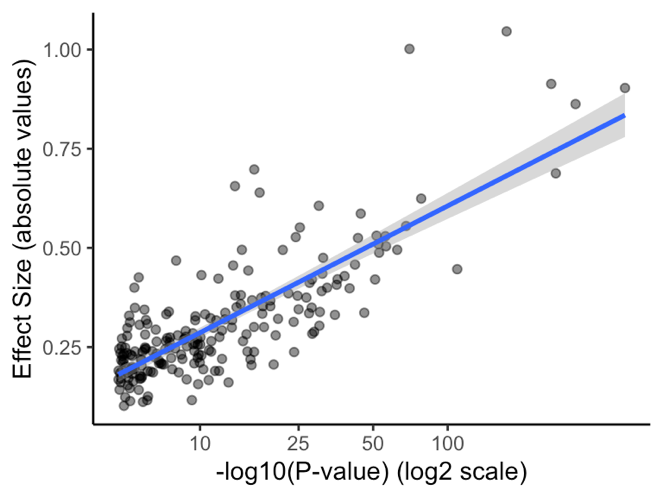

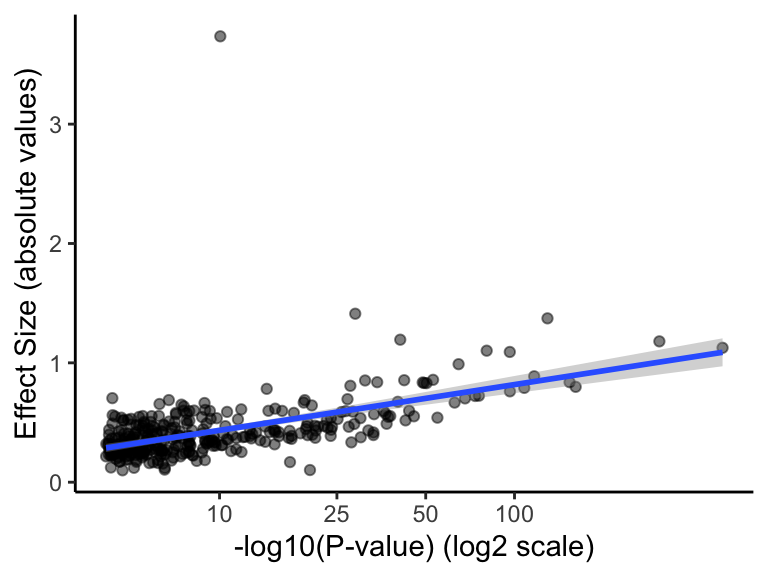

**Figure S.5. TE-eQTL Effect size as function of -Log10(*P*-value)** for LCL (left) and iPSC (right). Blue line: regression curve (linear model) and 95% confidence interval.

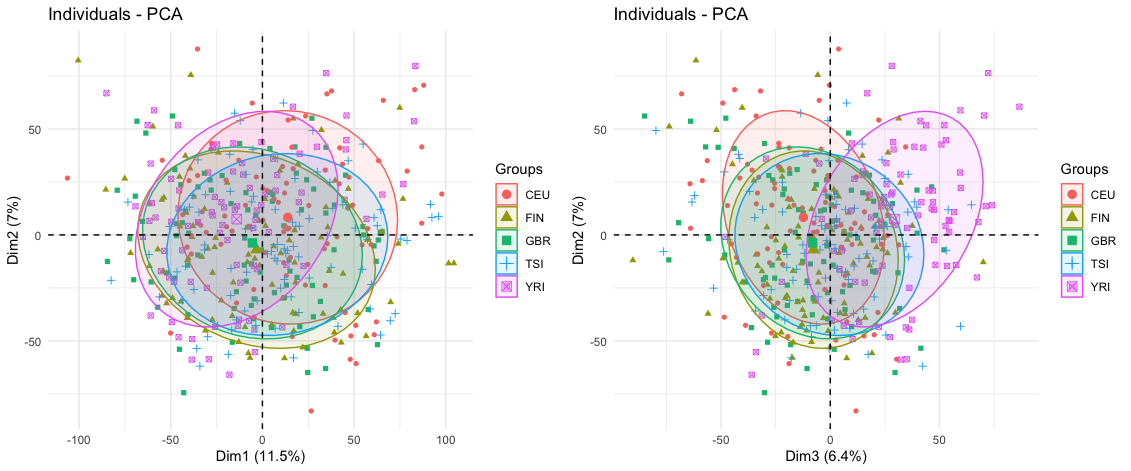

**Figure S.6. PCA of 444 LCLs based on normalized gene expression.** The structuration by continental populations (European ancestry populations [CEU, FIN, GBR, TSI] vs African [YRI]) only account for <= 6.4% of the total variance of the dataset.
